# *In utero* adeno-associated virus (AAV)-mediated gene delivery targeting sensory and supporting cells in the embryonic mouse inner ear

**DOI:** 10.1101/2024.05.07.592885

**Authors:** Carla Maria Barbosa-Spinola, Jacques Boutet de Monvel, Saaid Safieddine, Ghizlène Lahlou, Raphaël Etournay

**Affiliations:** Institut Pasteur, Université Paris Cité, Inserm, Institut de l’Audition, Paris, France; Centre National de la Recherche Scientifique, Paris, France; APHP Sorbonne Université, Département d’Oto-Rhino-Laryngologie, Unité Fonctionnelle Implants Auditifs,Groupe Hospitalo-Universitaire Pitié-Salpêtrière, Paris, France; Sorbonne Université, Collège Doctoral, Paris, France

## Abstract

In vivo gene delivery to tissues using adeno-associated vector (AAVs) has revolutionized the field of gene therapy. Yet, while sensorineural hearing loss is one of the most common sensory disorders worldwide, gene therapy applied to the human inner ear is still in its infancy. Recent advances in the development recombinant AAVs have significantly improved their cell tropism and transduction efficiency across diverse inner ear cell types to a level that renders this tool valuable for conditionally manipulating gene expression in the context of developmental biology studies of the mouse inner ear. Here, we describe a protocol for *in utero* micro-injection of AAVs into the embryonic inner ear, using the AAV-PHP.eB and AAV-DJ serotypes that respectively target the sensory hair cells and the supporting cells of the auditory sensory epithelium. We also aimed to standardize procedures for imaging acquisition and image analysis to foster research reproducibility and allow accurate comparisons between studies. We find that AAV-PHP.eB and AAV-DJ provide efficient and reliable tools for conditional gene expression targeting cochlear sensory and supporting cells in the mouse inner ear, from late embryonic stages on.

## Introduction

Understanding the molecular mechanisms underlying inner ear development is crucial not only for identifying the optimal time windows for the effective delivery of therapeutic molecules but also for improving the stepwise differentiation of stem cells into inner ear organoids. Notably understanding the role and the temporal dynamics of specific gene regulatory networks required for cochlear cell fate, patterning, and long-term survival will be instrumental to strengthen our view of the underlying pathophysiological processes, and therefore to optimize gene therapy outcomes. Such studies require finely tuned methods to achieve the highest levels of both efficiency and precision.

The emergence of inner ear gene therapy using recombinant adeno-associated viral vectors (AAVs) in clinical mouse models for human deafness has established AAVs as highly effective tools for conducting conditional genetic manipulations of specific cell types within the inner ear [1–3]. Several proof of principle studies have shown that the use AAVs for virus mediated gene transfer to the developing inner ear is an efficient and safe strategy for gene delivery [4,5], allowing the rescue of hearing function in several mouse models for inherited congenital human deafness mice presenting mutations causing profound deafness [6,7].

The cochlear sensory epithelium, the organ of Corti, harbors two types of sensory cells, the inner hair cells (IHCs) are the genuine sensory cells, which detect and transmit sound information to primary auditory afferent neurons. They are organized along a single row located most medially on the spiral-shaped longitudinal axis of the cochlea, the outer hair cells (OHCs) function as cochlear amplifiers of the sound-evoked vibrations of the organ. They are organized in three parallel rows (OHC1, OHC2, OHC3) located laterally to the IHC row. During cochlear embryonic development, cell differentiation proceeds in a longitudinal wave that initiates near the cochlear base and extends towards the apex [8]. The wave also progresses laterally from the medial to the lateral side, IHCs differentiating first at the cochlea base (from ∼E14.5 on), followed by the first row of OHCs (∼E15), and shortly after by the second (∼E15.5) and third (∼E16) OHC rows [9]. Sensory cells are interdigitated with supporting cells (SC) forming a highly regular mosaic arrangement of cells. For the needs of gene therapy, it is of great importance to optimize AAV serotypes to specifically and efficiently target the cell of interest such as sensory cells or supporting cells. Such tools would also be of great value for basic studies of the gene regulatory networks at play in the development of these cells, and for studies aiming to understand the function of a variety of genes implicated in cochlear physiology.

Numerous AAV serotypes have already been tested mostly in neonatal and to a lesser extent in mature mouse inner ear, exhibiting different transduction rates and expression levels in specific target cells and tissues [2,10,11]. The recombinant serotypes AAV-PHP.eB and AAV-DJ target the two major cochlear cell types, the sensory hair cells and the supporting cells, respectively, with remarkably high efficiency and specificity when administered at neonatal stages [12]. Both AAV-PHP.eB and AAV-DJ are recombinant serotypes whose capsids were engineered by an artificial evolution design, which results in the hybrid combination of eight different AAV serotypes: caprine, avian and bovine AAVs, and AAVs type 2, 4, 5, 8, and 9 [13–15]. The AAV-PHP.eB serotype derives from AAV9, and is identical to its parental capsid except for the addition of an extra heptamer amino-acid insertion in the capsid sequence, which confers to it the ability to cross the blood brain barrier [16–18]. Relatively low doses of AAV-PHP.eB (10^9 GC/ml) provide good transduction rates in both cochlear IHCs and OHCs [12]. The AAV-DJ serotype strongly transduces supporting cells when injected at neonatal stages in the mouse inner ear [12].

Unlike investigations involving *ex vivo* studies of embryonic inner ear development, where explanted tissues are cultured and incubated in the presence of AAVs [1], *in vivo* studies of cochlear development require the challenging task of performing *in utero* injection of AAVs to the inner ear, a technique that is still scarcely documented [5–7,19]. Importantly, there is currently no standardized method for evaluating precisely AAV transduction rates achieved in the cochlea. Considering their high capability to transduce cochlear cell types during postnatal stages, we sought to characterize the developmental time window during which the AAV-PHP.eB and AAV-DJ serotypes could effectively be used to manipulate gene expression in the two major cell types of the embryonic inner ear. We also characterized the temporal pattern of the transgene expression, with the aim of linking it to the cell maturation process along the cochlear length. In this protocol, we describe a comprehensive procedure for conducting *in utero* AAV injection, followed by image acquisition and analysis. Indeed, the mosaic pattern of the AAV transduction, where not all cell types of interest are transduced, requires careful consideration during the analysis and interpretation of the results. This difficulty can only be overcome by a multiscale approach that entails imaging of the entire tissue and performing various analyses down to the cellular scale. Our multiscale approach is designed to establish a standardized method for using AAV as a tool for conditional gene expression into specific cell types, using the cochlea as a model system.

## Materials and methods

The detailed in utero inner ear injection protocol described in this article is published on protocols.io, (DOI: dx.doi.org/10.17504/protocols.io.kqdg3x62pg25/v1). The injection setup is also shown in Supplementary Fig S1.

### Ethics declaration

All procedures used in this study complied with ethical regulations for animal research according to the European guide for the care and use of laboratory animals. The study received ethical approval from the ethics committee in animal experimentation of the Institute Pasteur under the APAFIS #31811–2021052715303576 (France).

### Animals

Timed-pregnant C57BL6 females were staged based on the day of conception, taking as reference (embryonic day E0.5) the morning at which the vaginal plug was observed. For each experiment, embryos from the same litter were all subjected to a unilateral injection of 1.2μl of viral solution. To estimate the precise stage of the embryos at the time of the AAV injection, hindlimbs of embryos of two independent litters at E13.5 from the same mouse strain were analyzed using the eMoss staging system [20]. The embryos were harvested also in the morning (Fig S5). With eMoss we found that the actual stage of C57BL6 embryos harvested in the morning of the 13th day after the night of mating corresponded to E13.5 delayed by 10h ± 4.1h.

### AAV vectors preparation

The AAV.PHP.eB::CAG-GFP [21] (2.6 X 10^13^GC/ml) was produced by Addgene, pAAV-CAG-GFP was a gift from Edward Boyden (Addgene viral prep # 37825-PHPeB; http://n2t.net/addgene:37825; RRID:Addgene_37825). The viral vectors were purified by iodixanol gradient ultracentrigation and titered by ddPCR in the manufacturer’s facility. AAV-DJ::CAG-GFP (1.3 X 10^13^ GC/ml) was purchased from Vector Biolabs (cat. No: 7078). The viral stock was purified by cesium chloride centrifugations from transfected HEK293 cells and the titer was determined by quantitative real time PCR by the manufacturer.

### Microscopy

Whole-mount stained cochleas were imaged with a W1-Yokogawa confocal spinning disk microscope (Gataca Systems) equipped with an inverted Nikon-Ti2 stand and with plan apochromat lenses. We used low magnification lenses for screening specimens: a dry 20x lens (NA 0.8) or a silicon oil immersion 25x lens (NA 1.05). For multiscale imaging of entire cochleas at subcellular resolution, we used either a water immersion 40x lens (NA 1.15), or a water immersion 60x lens (NA 1.2) in combination with a Photometrics 95B or BSI CMOS camera. The exposition times and percentage of fluorescence were configured for each cochlea using the ND-Scan software (Gataca Systems) for MetaMorph, and tile scans were acquired sequentially for each confocal plane in multiple stacks, for each channel. Stitching was done using Fiji [22,23].

Some cochlear areas of interest were also acquired with a Zeiss LSM900 Airyscan confocal microscope, using a 40x oil immersion objective (NA1.4). In this case, all configurations, as well as tiling and stitching of mosaic z-stack acquisitions, were done using the ZEN software.

### Image analysis

Transduced cells of each cochlea from each AAV-injected individual were manually selected and the (XY) image coordinates of their center were collected using the Fiji Multipoint selection tool. Each transduced hair cell was further categorized by its type (IHC, or OHC from row 1 to 3, the first OHC row being the closest to the IHC row). Similarly, each supporting cell type could be identified based on their location within the characteristic mosaic pattern of the cochlear neurosensory epithelium. The cochlear longitudinal axis was reconstructed by delineating a spiral spline running the length of the organ of Corti from base to apex, using the Fiji line segmenting tool. The image coordinates of the segmented cochlear spiral were saved to the ROI Manager tool, from which they could be easily exported for subsequent analysis, which was performed with Matlab (The Mathworks) using custom functions for the generation of longitudinal cochlear transduction profiles. The longitudinal coordinates of every transduced cell (defined as their distance from the base along the cochlear spiral) were first obtained by projecting their center onto the cochlear axis. A running average of these longitudinal coordinates was then computed using a Gaussian averaging window having a standard deviation of 250 µm (approximately 1/20th of the total cochlear axis length). We thus obtained a “cochleogram” giving the average number of transduced cells within the running window at any given point along the cochlear axis. The calculated cochleograms were finally converted to longitudinal transduction profiles giving the average fraction of transduced cells as a function of cochlear position (Fig 2 and 3). For this conversion, the following reference values for the longitudinal density of hair cells along the P0 cochlea were used: IHC: 11.8, OHC1: 13.7, OHC2: 13.9, OHC3: 14.1, in number of cells per 100µm, estimated in 3 cochleas. Statistical analyses were performed in Matlab or in Excel, using the Kolmogorov-Smirnov test for comparison of longitudinal cochlear profiles, or Welch’s t-test for comparison of mean values.

## Results

### AAV-PHP.eB and AAV-DJ cell tropism in neonatal cochleas

Various factors, such as the manufacturer, purification process, delivery route and the developmental stage at which the in vivo injection is performed, can influence the tropism of a specific AAV serotype. Therefore, we first aimed to validate the tropism of the AAV-PHP.eB and AAV-DJ serotypes under our specific experimental conditions. To do so, we delivered 1.2µl of either one of these serotypes at a titer of ∼10^13^ genome copies per ml (GC/ml) through the round window membrane of the left inner ear in neonatal P3 C57BL6 mice [24,25]. Both AAV-PHP.eB::CAG-GFP and AAV-DJ::CAG-GFP express a GFP reporter gene under the control of the ubiquitous CAG promotor. Both left and right cochleas of each individual were harvested 5 days post-injection, at postnatal day 8 (P8), and whole-mount preparations of the cochlear epithelium were stained for GFP and F-actin, followed by imaging using confocal microscopy. The cochleas injected with AAV-PHP.eB::CAG-GFP presented high transduction rates of both IHC and OHC. To assess the local transduction rate profiles of AAV-PHB.eB::GFP of the different sensory cell types in relation to their position along the longitudinal axis of the cochlea, we manually collected the positions (X,Y image coordinates) of the transduced cells classified according to cell type (IHC, OHC1, OHC2, OHC3), using the Fiji multipoint selection tool for one individual. From these coordinates, we obtained longitudinal density profiles (or ‘cochleograms’, in number of transduced cells per µm) by counting the number of transduced cells from base to apex along the cochlea. These histograms were then smoothed by a Gaussian running window and converted into the transduction rate profiles shown in (Fig 1A-D). IHCs were overall the most transduced, with transduction rates gradually increasing from 20-80% at the cochlear base to 80-90% in the mid region and reaching ∼100% at the cochlear apex (Fig 1A-D). OHCs were also more transduced at the mid and apical regions (40-95%) than at the base (30-70%), with some substantial local variations throughout the cochlea. These results are consistent with those obtained by Hu et al. In the contralateral cochleas (non-injected ear), we found that a few sensory cells (<20% of IHCs and <10% of OHCs) were transduced by the virus (Fig S2A). These results are consistent with earlier studies in the neonatal mouse cochlea [26,27], in which the injection of AAV or other agent through the round window in one cochlea was reported to reach the contralateral cochlea (presumably through the cochlear aqueduct) and to transduce cells at variable rates.

**Fig 1.**
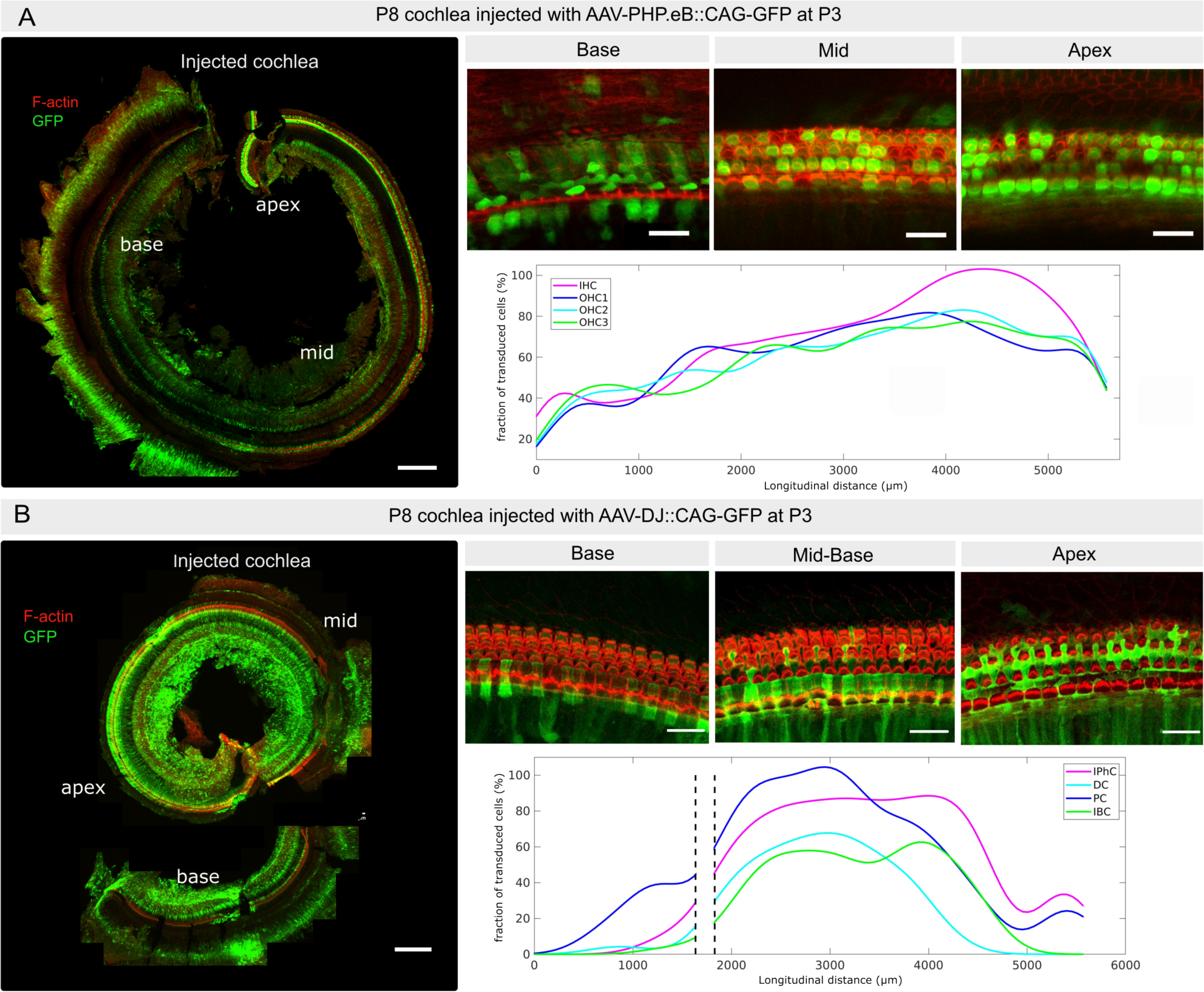
Inner ear transduction efficiency of AAV-PHP.eB and AAV-DJ administered at the neonatal stage. (A) P8 cochlea injected at P3 with AAV-PHP.eB through the round window membrane. The image on the left shows F-actin (red) and transduced cells expressing GFP (green) in maximum projection in the same cochlea. Upper panels on the right show representative close-ups of the sensory epithelium at the cochlear base, mid, and apex; the graph below shows the corresponding longitudinal transduction profile (fraction of transduced sensory cells as a function of position) along the cochlear axis, the latter being oriented from base, x = 0, to apex, x ≃ 5700 µm). IHC: Inner hair cells; OHC1-3: Outer hair cells from row 1-3. (B) P8 cochlea injected at P3 with AAV-DJ through the round window membrane. The image on the left shows F-actin (red) and transduced cells expressing GFP (green), in maximum projection in the same cochlea that was cut into two pieces. Panels on the right show representative close-ups of the sensory epithelium (base, mid and apex); the corresponding longitudinal transduction profile (shown below) was obtained by combining the partial profiles obtained in the respective cochlear fragments. Note that AAV-DJ also transduces the inner sulcus and mesenchymal cells. IPhC: Phalangeal cells; DC: Deiter cells; PC: pillar cells; IBC: inner border cells. Scale bar 200 µm.

The transduction rate of AAV-DJ::CAG-GFP in the mid-apex supporting cells exceeded that observed at the base (Fig 1E-H). The supporting cell types showing a sizeable rate of transduction included the inner phalangeal cells, Deiters’ cells, pillar cells, and the inner border cells. Pillar cells exhibited the highest transduction rates, increasing from the base (70-80%) to reach a maximum in the mid cochlear region (90-100 %), and plummeting in the mid-apical regions (70-10%). Inner phalangeal cells were also transduced at a high rate (70-80%), which remained roughly constant along most of the cochlear length, while Deiters’ cells and inner border cells displayed comparable longitudinal transduction profiles with a peak in the mid cochlear region, but with lower rates (40-60%). Again, with the injection procedure adopted, the contralateral cochleae exhibited only few detectable transduced cells by AAV-DJ::CAG-GFP (Fig S2B).

In addition to their main targets, both the AAV-PHB.eB and the AAV-DJ viruses transduced various other cell types within the cochlea, including inner sulcus and interdental cells, mesenchymal cells, and cells within Reissner’s membrane. These results are overall consistent with the previous study by Hu et al [12]; validating the quality of the AAV preparations used, and confirming that a diffusion of the AAV solution to the contralateral ear is possible at the neonatal stage.

### Dye unilaterally injected into the embryonic inner ear diffuses to the contralateral ear

The high transduction rates of the AAV-PHP.eB and AAV-DJ serotypes observed at postnatal stages raised hope regarding their potential use for implementing conditional gene expression within sensory and supporting cells during cochlear development. This would provide a key tool for example, to disentangle the mechanisms underlying autonomous versus non-autonomous cell differentiation, shedding light on their respective influence on patterning during the development of the cochlear sensory epithelium. One successful example has been achieved by Wu’s group in *ex vivo* experiments conducted on cultured vestibular utricle explants, using the AAV2.7m8 serotype [1,28]. Therefore, we sought to investigate the *in vivo* effectiveness of recombinant AAV-PHB.eB and AAV-DJ vectors during embryonic stages.

Unlike postnatal injections, *in utero* injections are challenging due to the difficult access to the inner ear of the embryos, making it difficult to precisely control the injection site. We defined a specific inner ear location based on morphological landmarks observed through the stereomicroscope. From E13.5 onwards, the external ear is clearly visible through the uterine wall, being positioned at the center of a region limited by the cephalic vein (located in the front) and its anterior and posterior branches (located above and below) (Fig 2A). After the needle was positioned closely to this landmark, injection was performed towards the labyrinth with the hand stably fixed to minimize any variations in the angle between the needle and the skin surface around the inner ear site. To fine-tune the injection procedure, we conducted pilot experiments where the Fast-Green dye was injected into E13.5 embryos. Those embryos were then harvested around 10 min after injection. We found that in utero injection resulted in the rapid transfer of the dye up to the contralateral (non-injected) inner ear (Fig 2A-C). The inner ear fluids are known to communicate with the cerebrospinal fluid (CSF) through in cochlear aqueduct [29]. These results are consistent with the notion that a rapid spread of AAV vectors should occur from the injected ear to the contralateral one, most likely through the CSF. Due to surgical constraints, and to limit the duration of the procedure, we made no attempt to distinguish which of the two ears (left or right) had been injected.

**Fig 2.**
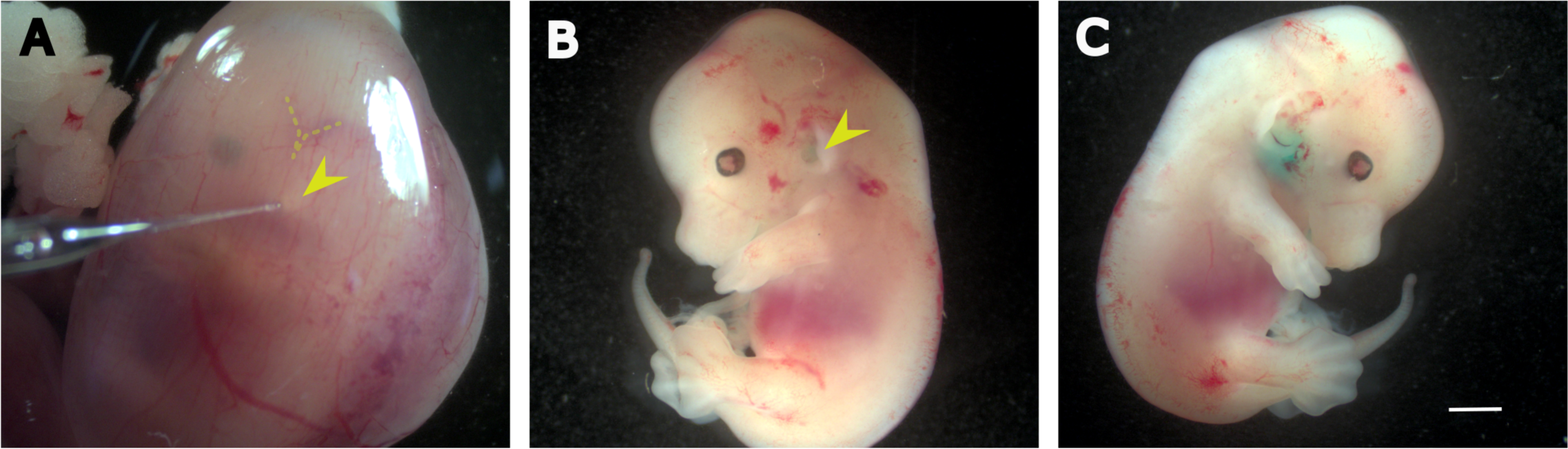
Inner ear diffusion of FastGreen administered at embryonic stage E13.5. (A) Photo of an E13.5 embryo inside the uterine horn. Notice the capillary pointing towards the external ear, located bellow the ‘V’-shaped veins corresponding to the transverse and sigmoid sinus veins. (B-C) Micrographs of E13 mouse embryos after the inner ears were injected with FastGreen. (B) The injected (left) ear. (C) The contralateral ear, which shows a higher concentration of dye. Scale bar 1 mm.

### AAV-PHB.eB transduces cochlear hair cells at early embryonic stages

We subsequently conducted intra-uterine injections of AAV-PHB.eB::GFP. All embryos of the same dam received unilateral injection either on E13.5, when the cochlear sensory epithelium cells initiate their differentiation into sensory and supporting cells at the cochlear base, or one day later, on E14.5 to examine a possible link between the transduction efficiency and the developmental stage. Both the right and left inner ears were subsequently collected at birth, fixed, and subjected to F-actin staining for analysis. We found that the unilateral injection of AAV-PHP.eB::CAG-GFP at either E13.5 or E14.5 resulted in robust transduction of hair cells of both the left and right cochleas at P0, with transduction rates that were highest in the basal cochlear region (Fig 3A-B). In addition, all cochleas injected at E13.5 or at E14.5 presented GFP-positive filaments, morphologically identified as the dendrites of type I afferent neurons facing the synaptic zone of the IHC row along the longitudinal axis (Fig S3A,B). This was expected, as AAV-PHP.eB is known to have a capacity for retrograde and anterograde transport across the axons of the central nervous system [30]. In cochleas where some mesenchyme was preserved, we also observed a strong GFP signal emanating from cells positioned medially to the sensory epithelium (Fig S3B), and the GFP-labeled neuronal fibers, identified as the spiral ganglion neurons based on their location and morphology. Of note, these cell types were not transduced when this serotype was injected at postnatal stages (not shown).

**Fig 3.**
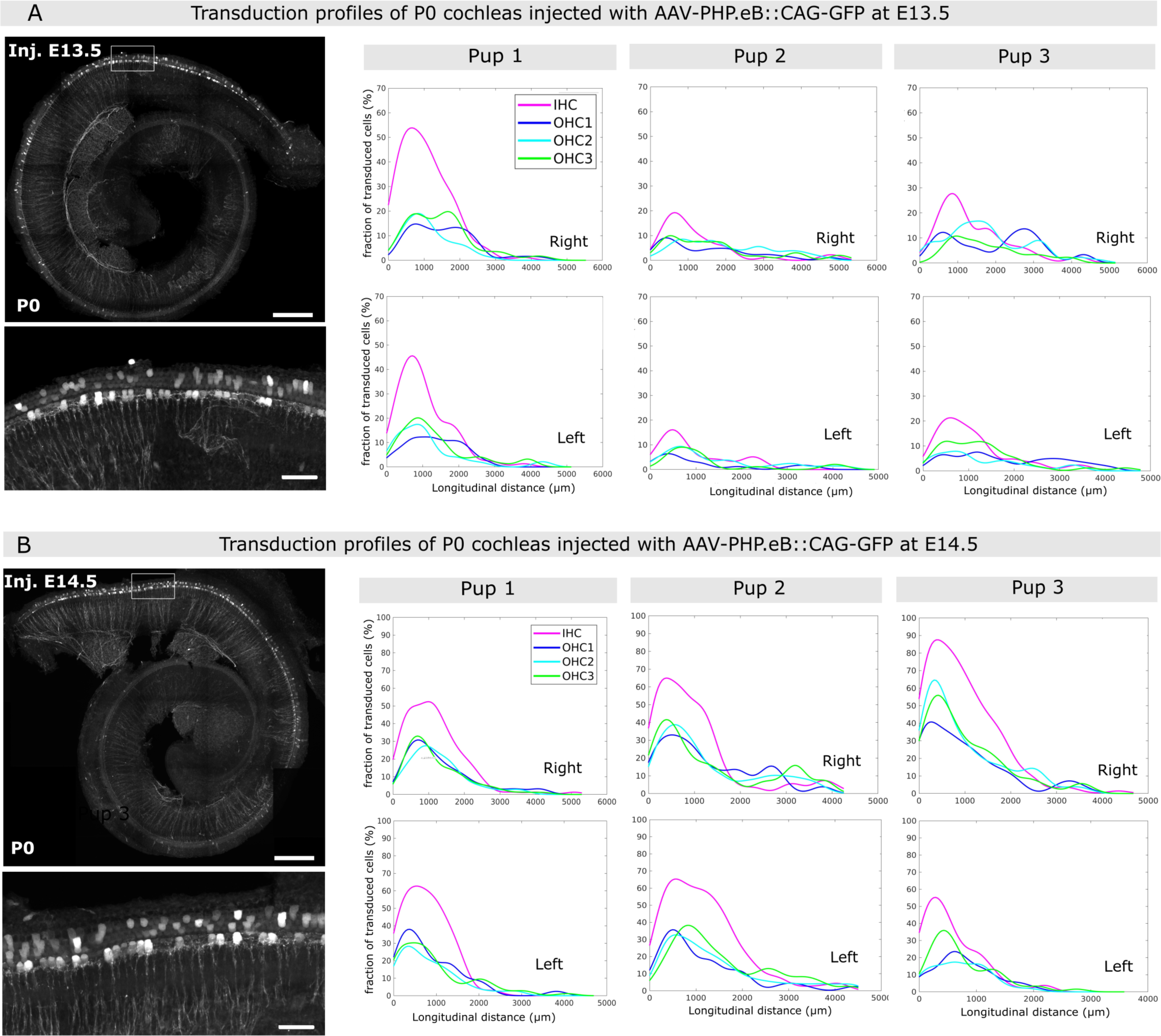
Inner ear transduction efficiency of AAV-PHP.eB administered at embryonic stages E13.5 and E14.5. (A-B) Whole-mount P0 cochleas injected with AAV-PHB.eB::GFP at E13.5 (A) and at E14.5 (B). Images on the left show stitched and projected views of the whole-mount cochleas imaged by mosaic confocal microscopy. Inset images below show the transduced sensory cells in more detail. Scale bars in overview images, 200 µm, in the insets, 25 µm. The graphs on the right show representative longitudinal transduction rate profiles measured in the right and left cochleas of 3 pups injected at E13.5 (A), and 3 pups injected at E14.5 (B). See also Supplementary Fig. 3, which contains additional transduction rate profiles for 3 pups injected at E13.5 and 3 pups injected at E14.5. IHC, inner hair cells, OHC1, OHC2, and OHC3, first, second and third row of outer hair cells, respectively.

To determine the viral transduction rates, we conducted a cell count for each sensory cell type (IHC, OHC1, OHC2, OHC3). Representative transduction rate profiles deduced from these counts are shown for both stages of injection in Fig 3C-D and FigS3C-E. The total cell counts (including all sensory cell types) are reported in Table 1 for injections at E13.5 and in Table 2 for those at E14.5.

**Table 1.** Total numbers of GFP-positive hair cells per individual: AAV-PHP.eB injected at E13.5.

| Individual | Cell number (left cochlea) | Cell number (right cochlea) |
| --- | --- | --- |
| **Pup1** | 94 | 87 |
| **Pup 2** | 125 | 74 |
| **Pup 3** | 118 | 186 |
| **Pup 4** | 259 | 204 |
| **Pup 5 (Fig S3C)** | 166 | 480 |
| **Pup 6 (Fig S3C)** | 218 | 180 |
| **Pup 7 (Fig S3D)** | 536 | 854 |

**Table 2.** Total numbers of GFP-positive hair cells per individual: AAV-PHP.eB injected at E14.5.

| Individual | Cell number (left cochlea) | Cell number |
| --- | --- | --- |
| **Pup 1** | 312 | 326 |
| **Pup2** | 396 | 401 |
| **Pup3** | 190 | 520 |
| **Pup 4** | 186 | 128 |
| Pup 5 (Fig S3E) | 49 | 29 |
| Pup 6 (Fig S3E ) | 720 | unavailable |

Qualitatively, we obtained similar transduction profiles on P0 cochleas for injections conducted at E13.5 and E14.5. In particular, for inner ear injections conducted at E13.5, the regional transduction rate displayed a peak value at the cochlear base for both IHCs and OHCs (ranging 10-55% for IHCs, and 7-35% for OHCs), then gradually decreased towards the cochlear mid, and vanished at the mid-apical region. At the cochlear base, IHCs were transduced on average twice as frequently as OHCs (peak transduction rate averaging 25.3 ± 1.2% for IHCs, and 12.7 ± 0.6% for OHCs, p = 0.012, t-test, 12 cochleas from 6 pups). At the cochlear mid region, the transduction rates of the two sensory cell type became similar. In one individual from the litter that received injection on E13.5, we observed a different transduction pattern where both IHCs and OHCs displayed unusually high transduction rates (reaching ∼65% for IHCs and 55% for OHCs), with a reversed trend of higher transduction rates at the cochlear apex compared to the base (Fig S3D). Except for this particular individual that was considered an outlier, the transduction rates generally observed after an injection conducted at E13.5 were ∼3 times lower than those observed after a neonatal injection (p<0.001, t-test; see Fig 1,3 and S3). For ear injections conducted at E14.5, the peak transduction rates measured in these cochleas were also located at the cochlear base and were also on average higher in the IHCs (52.1±1.5%) than in the OHCs (29±1.8%, p = 0.036, t-test), but they were about twice higher than those from the pups injected at E13.5 (p < 0.015, t-test, 11 cochleas from 6 pups). Since we injected the same volume of the same viral preparation for all pups, our results are consistent with a temporal transduction pattern of the sensory cells that reflects their state of differentiation. Yet, the variability of these profiles and the atypical profile observed (Fig S3D) suggest that other factors than the maturation of the cochlear epithelium also likely contribute.

Despite the unilateral injections and a substantial variability between pups, both cochleas of each pup exhibited comparable transduction profiles, both in terms of the average number of transduced hair cells in various cochlear segments (t-test, p>0.05) and in the shape of these profiles (Kolmogorov-Smirnov test, p>0.05). This feature is not too surprising in view of the results obtained with the Fast-Green dye injection (Fig 2). We conclude from these results that a robust hair cell transduction could be obtained at both stages of injection, the transduction rate being lower for injections performed at E13.5 than at E14.5.

### Temporal evolution and variability of AAV-PHB.eB transduction rates across the post-injection period

To gain a deeper understanding of the dynamics of AAV-PHP.eB::GFP transduction in the maturing cochlear epithelium, cochleas injected on E13.5 were harvested for analysis at 48h and 72h post-injection, *i.e* on E15.5 and E16.5, respectively (Fig 4A). At the 48h post-injection stage, most of the GFP-expressing cells were located underneath the surface of the cochlear epithelium. In three individuals analyzed (5 cochleas, one cochlea having been damaged), only scarce GFP-positive hair cells were observed within the cochlear epithelium (Fig 4A, upper panels and Fig S4A). At the 72h post-injection stage, a notably greater number of GFP-positive were found in the organ of Corti (Fig 4A, lower panels). The right and left cochleas of each mouse pup exhibited comparable transduction rates (in the range 3-15% for IHCs and 5-20% for OHCs), but there was variability in the transduction profiles along the cochlear longitudinal axis among the three individuals. In two individuals (pups 3 and 4 in Fig 4B), the sensory cells displayed significant transduction rates at the cochlear base, mirroring the results seen in the P0 cochleas that were injected at E13.5 (see Fig 3E above). In the third individual (pup 6 in Fig S4B), we observed an overall higher transduction rate, with a maximum at the cochlear mid-apical region (reaching ∼10% of IHCs and ∼20% of OHCs). These observations again indicate that the transduction rate may not only reflect the maturity state of the cells but is also likely influenced by other (e.g. physical and geometrical) factors conditioning the injection. Variations in these factors, especially in the angle of approach of the needle, may have modified the injection route, leading to different spreading patterns of viral particles (∼3750 kDa) across the inner ear, although the same concentration (∼10^13^ GC/ml), volume (1.2 µl), and flow speed (0.6 µl/min) were consistently used for all injections. In addition, the overall transduction rate increased over time post-injection (2 days, 3 days, or 6 days), which may correspond to the time required for viral entry into the cells and the subsequent transduction.

**Figure 4.**
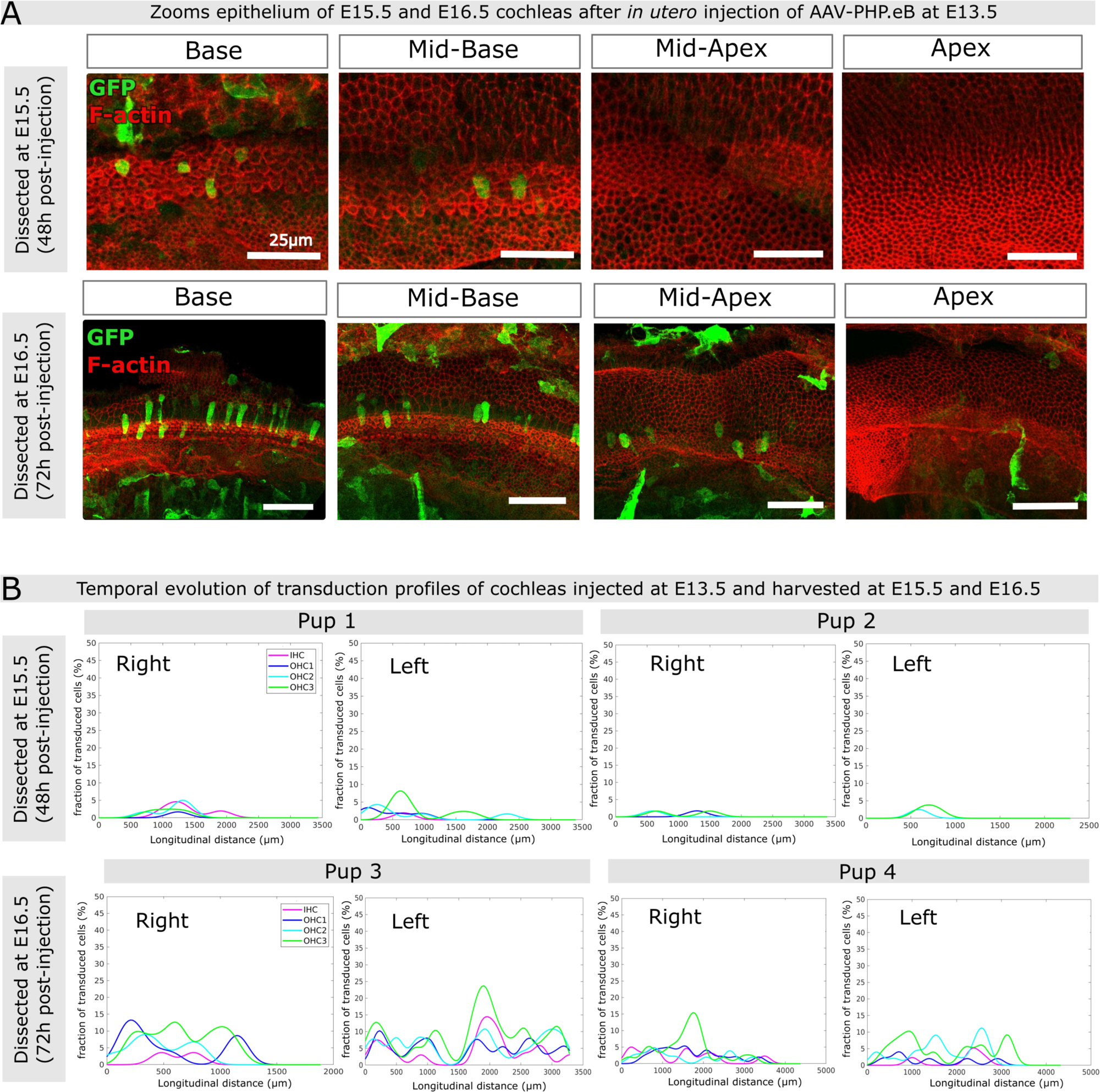
Inner ear transduction efficiency of AAV-PHP.eB administered at embryonic stages E15.5 and E16.5. (A) Confocal views (maximum intensity projections) of the cochlear epithelium of mice subjected to unilateral injection of AAV-PHB.eB::GFP at E13.5, and imaged at E15.5 (upper panels) or at E16.5 (lower panels). Panels from left to right show different cochlear locations from base to apex. Staining for F-actin (red) and GFP (green). Scale bars 25 µm (upper panels) and 50 µm (lower panels). (B) Longitudinal transduction rate profiles observed in cochleas dissected at either E15.5 (upper panels) or E16.5 (lower panels) after unilateral injection of AAV-PHP.eB at E13.5. IHC, inner hair cells, OHC1, OHC2, and OHC3, first, second and third row of outer hair cells, respectively.

### AAV-DJ transduces cochlear supporting cells at late embryonic stages

Six embryos from the same dam were injected at E13.5, with the same viral preparation that we used for P3 injections of AAV-DJ::CAG-GFP. Upon dissection at birth, most of the cochleas presented no GFP-positive cells. Only one cochlea out of six contained a few GFP-positive cells identified as sensory cells based on their morphology and location within the apex and mid regions of the organ of Corti (Fig 5A). Therefore, the unilateral delivery of AAV-DJ::CAG-GFP into the inner ear at E13.5 did not yield appreciable transduction of supporting cells, or any other cell type in cochleas analyzed at P0. Considering that the AAV-DJ efficiently and specifically transduces supporting cells during neonatal stages, this suggested that the receptors to this serotype might only be expressed at late embryonic stages as the supporting cells differentiate. To test this hypothesis, we injected the same volume of this viral preparation to embryos at E14.5, E15.5 and E16.5 and subsequently analyzed the cochleas at P0 (Fig 5A,B). Delivery of AAV-DJ::CAG-GFP at E14.5 to five embryos from the same litter, once again did not lead to exploitable transduction of the cochlear cells. However, AAV-DJ::CAG-GFP delivery at E15.5 and E16.6 resulted in consistent and comparable transduction profiles of the supporting cells (Fig 5C), as well as of the inner sulcus cells, from the basal region of the epithelium (Fig 5B) in both the left and right cochleas. The transduction rates in P0 cochleas subjected to injection at either E15.5 (2 cochleas from 1 pup) or E16.5 (5 cochleas from 3 pups) were similar, with peak values ranging 10-50% for inner phalangeal cells, pillar cells, and inner border cells. The Deiters’s cells remained scarcely transduced in these cochleas, however, with peak transduction rates not exceeding 5% (Fig. 5C). Collectively, these results reveal a transduction pattern that progresses from the cochlear base to the apex, mirroring the differentiation dynamics of the cochlear epithelium. This is consistent with a dependency of the AAV-DJ::CAG-GFP transduction efficiency on the differentiation state of the supporting cells. Yet, other factors, particularly the variable conditions of injections combined with the complex geometry of the inner ear labyrinth, likely contributed to the observed variability of the transduction profiles.

**Fig 5.**
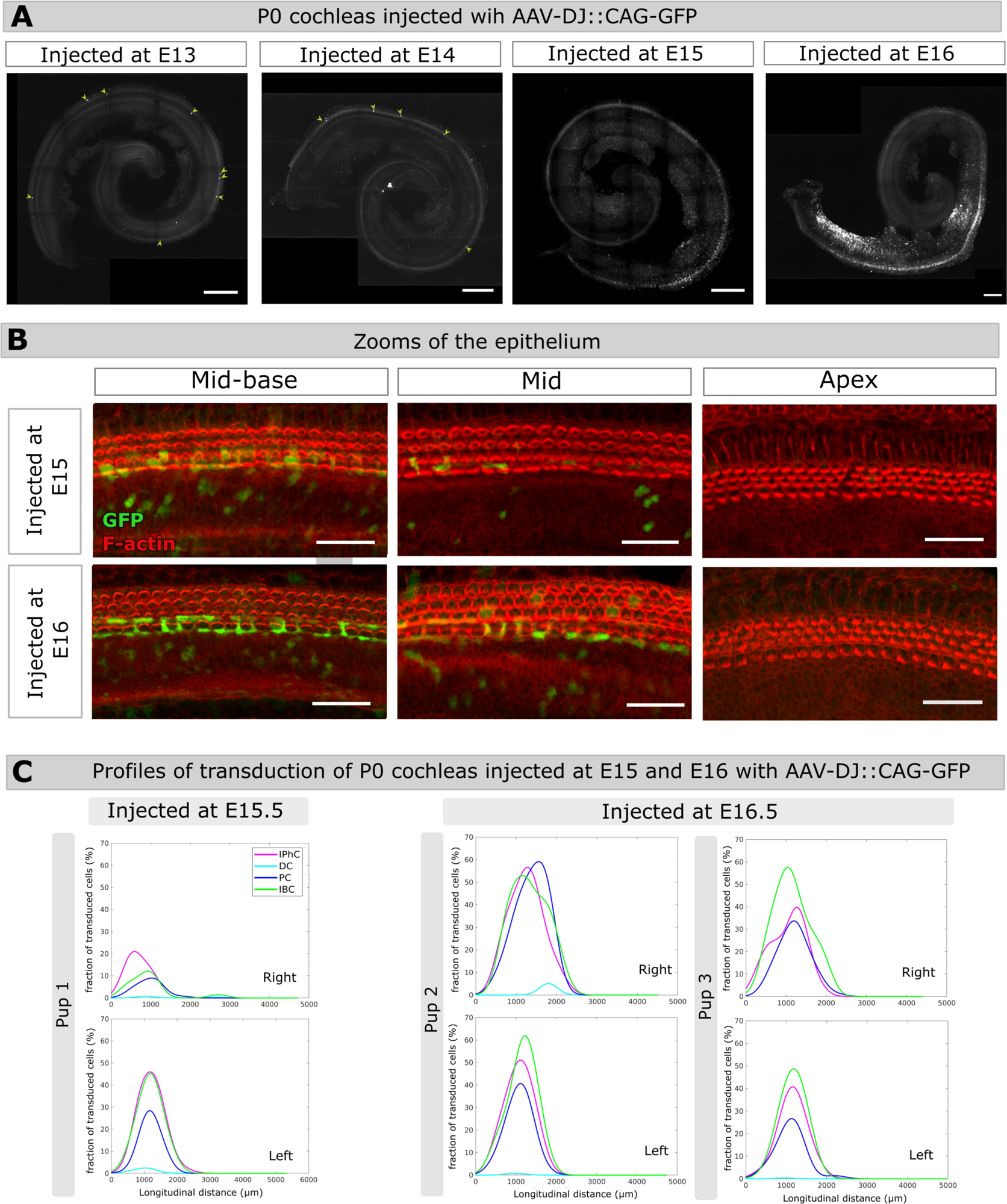
Inner ear transduction efficiency of AAV-DJ administered at embryonic stages. (A) Whole-mount P0 cochleas injected with 1.2µl AAV-DJ at E13.5, E14.5, E15.5, E16.5. Scale bar 200 µm. (B) Close-ups of the mid-basal, mid, and apical regions of P0 cochleas injected either at E15.5 (upper panels), or at E16.5 (lower panels). Scale bar 25 µm. (C) Longitudinal transduction rate profiles measured in the right and left cochleas of three different pups injected at E15.5 (pup 1) or at E16.5 (pups 2 and 3). IphC, phalangeal cells, DC, Deiter cells, PC, pillar cells, IBC, inner border cells.

## Discussion

This study represents to our knowledge the first quantitative characterization of AAV transduction rate profiles along the full length of the mouse cochlea during and after fetal development. Based on our findings, we conclude that the AAV-PHP.eB and AAV-DJ serotypes effectively transduce auditory sensory cells and supporting cells, respectively, at embryonic stages covering the period of cochlear extension and differentiation. Importantly, we confirmed that following a single unilateral injection of either serotype to the inner ear during fetal development, the therapeutic agent can readily diffuse to the contralateral inner ear. Our quantification shows moreover that under such conditions the longitudinal AAV transduction profiles observed in the left and right cochleas are not significantly different.

Our quantification also revealed that with just one exception, in all of the P0 cochleas subjected to AAV-PHP.eB injection at E13.5 or E14.5, the longitudinal transduction profiles of hair cells displayed higher transduction rates at the cochlear base compared to the apex. This clearly supports a pattern of AAV transduction correlating with the basal-to-apical wave of differentiation of sensory cells. However, the variability observed in our experiments, and the reversed transduction profile seen in one of the pups injected at E13.5, indicate that other factors than sensory cell maturation, such as asymmetries in the accessibility of the virus to these cells depending on the site of injection also likely influence this pattern.

Most in utero AAV gene delivery studies so far have focused primarily on sensory hair cells, so that comparatively little is known about the differentiation of supporting cells. Our results provide proof of principle for using AAV-DJ as a gene delivery tool effectively targeting supporting cells from E15.5 on, the earliest stage at which these cells seem to become competent for transduction.

Our data also suggest that it takes around 2 days post-injection to detect GFP expression in the cochlear epithelium. These results are in line with those of Tona et al. who were able to induce a reversal of hair bundle polarity in the sensory cells of the utricle approximately 2 days after incubation of the AAV-Emx2-tdT in organotypic culture [1]. Considering the expected ∼48h delay for the transgene to be expressed in the target cells, we would not expect this protocol to be effective to manipulate gene expression before E15.5. It can thus be anticipated that intended effects brought by the exogenous expression of the transgene may not become manifest at the cellular level until approximately 2 days have elapsed. Therefore, the AAV-PHP.eB and AAV-DJ serotypes appear well-suited for investigating late embryonic or neonatal inner ear development in the mouse. Notably, in utero injections into the mouse inner ear become considerably more challenging at stages prior to E13.5 (E12 is the earliest stage at which it could be successfully achieved so far, the mouse otocyst being formed at ∼E9.5 [31]).

A remaining limitation for in utero gene therapy is that undesired cell types, such as spiral ganglion cells, cells of the Reissner’s membrane, and also some brain regions, can be reached by the through the cerebrospinal fluid, and potentially transduced at notable levels. Indeed, we found that AAV-PHP.eB, when delivered at either E13.5 or E14.5, presents a high tropism not only to the sensory hair cells, but also to the spiral ganglion neurons and mesenchymal cells. This limitation could potentially be overcome through the use of specific promoters suitable for the embryonic period. Another limitation is that the competence of target cells for transduction evolves with the developmental stage. This is typically the case for the supporting cells, so that the AAV-DJ serotype does not transduce these cells unless injected after E15.5.

Therefore, caution should be advised when interpreting the the observed AAV transduction rates observed in hair cells or supporting cells with regard to their developmental stage.

The multiscale approach that we used in this study, which enables the quantification of AAV transduction rates along the full cochlear length, holds promise for conducting conditional gain and loss of function studies of specific cell types of within the organ of Corti during late embryonic or neonatal stages, particularly when using AAV-DJ and AAV-PHP.eB serotypes. Other serotypes, such as rAAV2/Anc80L65 [5] or rAAV2/1 [4,6,7] could also potentially be considered for late embryonic studies since in utero injection at E12.5 gives rise to a high transduction rate of sensory cells in P7 and P0 cochlea respectively. This approach should also help fostering reproducible research and further comparisons between studies, notably in the context of current joint efforts to synergize gene therapy with developmental biology studies of the inner ear.

## Supporting Information

**S1 File. In utero inner ear AAV injection protocol**

**S2 File. ARRIVE guidelines 2.0: author checklist.**

## Supporting information

https://www.protocols.io/view/in-utero-inner-ear-aav-injection-kqdg3x62pg25/v1

## Acknowledgments

We thank Maia Brünstein of the Hearing Institute Bioimaging Core Facility - C2RT for sharing expertise on light microscopy. We thank the staff of the Hearing Institute Animal Phenotyping Core Facility - C2RA for their help with mouse husbandry and ethical support. We thank Céline Trébeau for sharing expertise in image analysis, and Kelly Burkhead for helping with image analysis. We thank Charlotte Calvet, Marie Giorgi, Marie-José Lecomte for useful discussions about AAVs. We thank Mireille Montcouquiol, Christine Vesque, Sandrine Etienne-Manneville for insightful discussions on the project.

## Metadata

### Competing interests

The authors declare no competing interests.

### Associated content

DOI: dx.doi.org/10.17504/protocols.io.kqdg3x62pg25/v1

---

**Fig S1.**
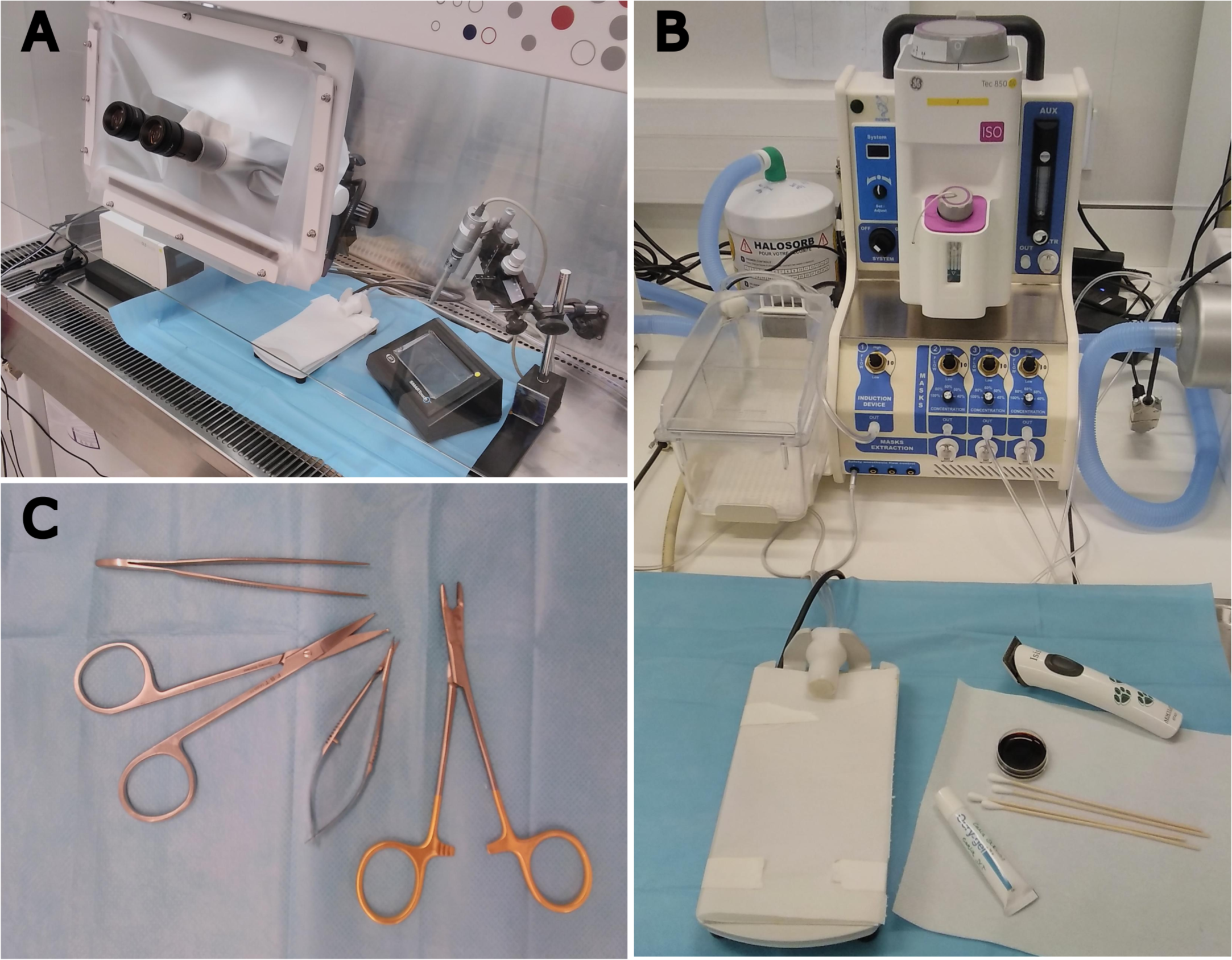
Surgery and injection materials. (A) Safety cabinet equipped with a binocular in the operatory area. (B) The preparatory area equipped with an anesthesia set up. In this area the pregnant dam is shaved in the stomach area after which an antiseptic solution of vetedine is applied. A solution of ocrygel is applied to both eyes. (C) Graefe forceps, ball-tipped scissors, microfine scissors as well as a needle holder used for the ventral laparotomy, as well as suturing.

**Fig S2.**
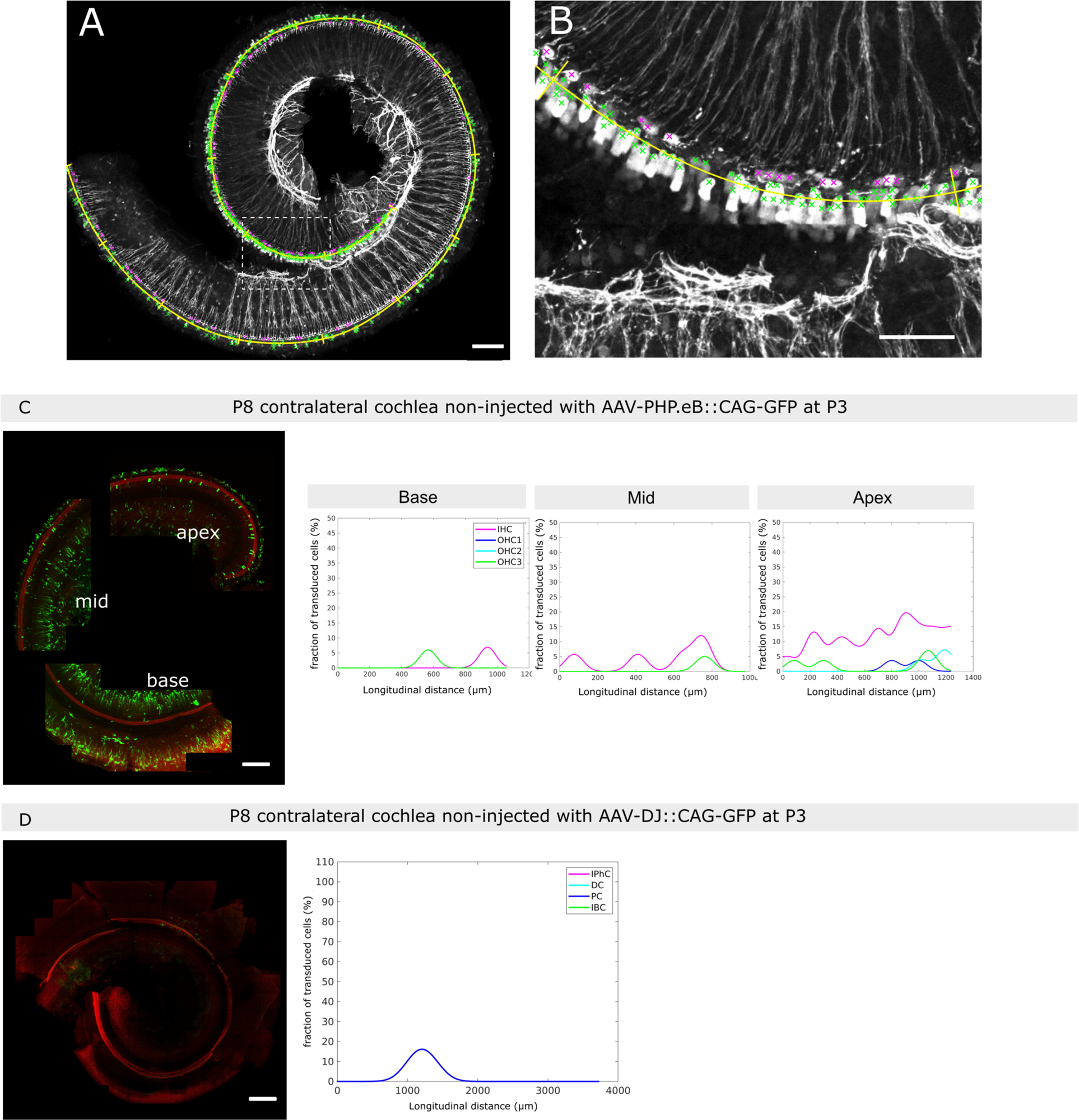
Quantification method to count transduced cells. (A) Confocal maximum projection image of a P0 cochlea injected with AAV-PHP.eB::CAG-GFP on E13.5. Sensory cells expressing GFP are in white. The longitudinal cochlear axis selected is shown (yellow spiral curve), together with the selected transduced cells with crosses in magenta for IHCs and in green OHCs. (B) Detail on the area delineated by the tireted rectangle in (A). (C,D) P8 contraleral cochleas from P3 injected pups. Confocal maximum intensity projection images of right cochleas extracted from pups injected unilaterally in the left ear with AAV-PHP.eB::CAG-GFP in (C) and AAV-DJ::CAG-GFP in (D). The graphs on the right of the images show the longitudinal transduction rate profiles estimated for each of the imaged cochlear fragments. Scale bars 100 µm.

**Fig S3.**
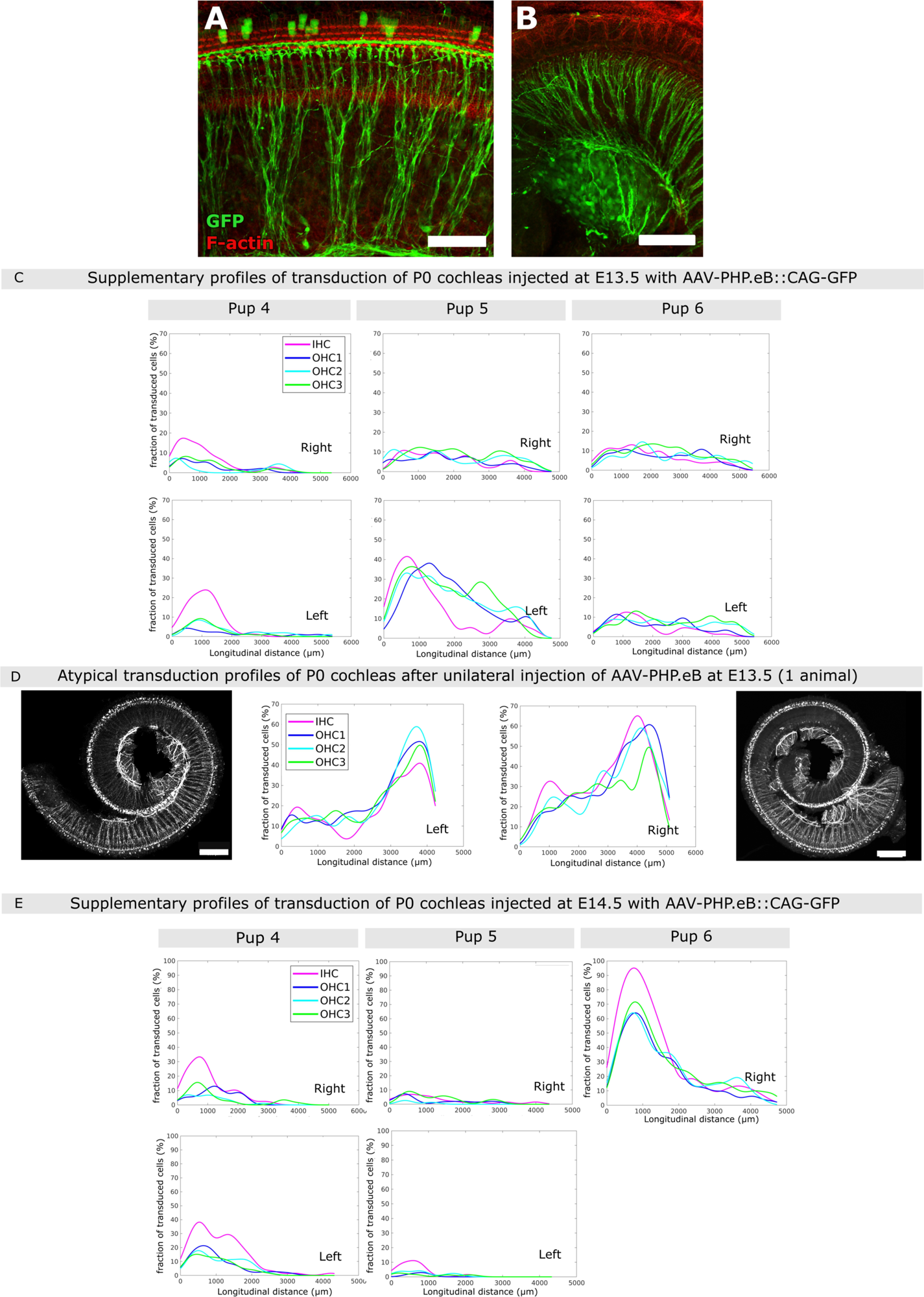
Cochlear cell transduction by AAV-PHP.eB::GFP. (A,B) Auditory nerve fibers (A) and ganglion-like cells (B) expressing GFP, in P0 cochlea injected at E13.5. Scale bar 100 µm. (C) Additional representative graphs of the longitudinal transduction profiles measured in the right and left cochleas of 3 individual pups injected at E13.5, and 3 pups injected at E14.5. (D) Longitudinal transduction profiles measured in the right and left cochleas of an atypical individual showing a reversed transduction profile. (E) Longitudinal transduction rate profiles observed in P0 cochleas from three additional pups after unilateral injection of AAV-PHP.eB at E14.5.

**Fig S4.**
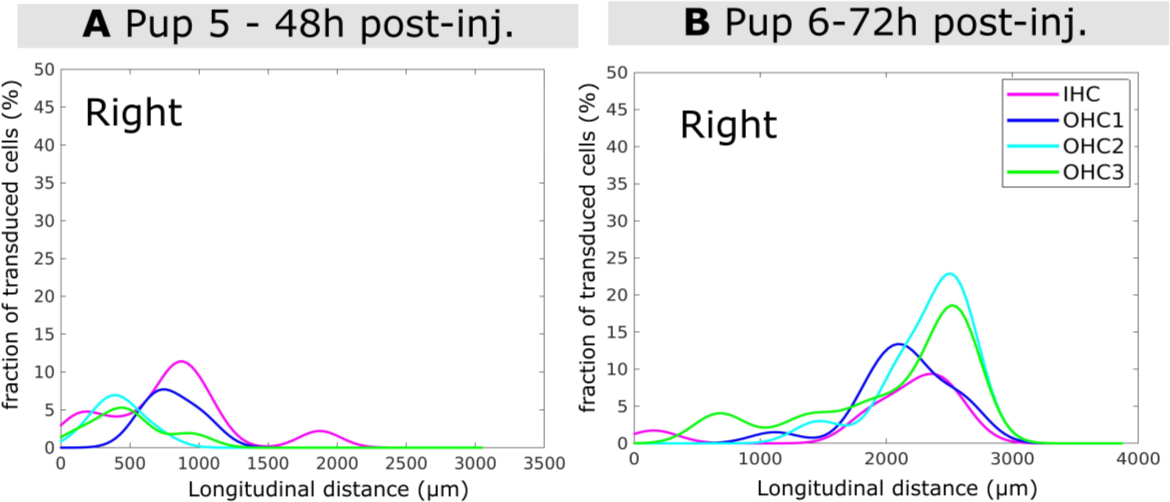
Transduction profiles of E15.5 and E16.5 cochleas injected at E13.5. Longitudinal transduction rate profiles observed in the right cochleas from two additional pups dissected 48 hours (at E15.5) (A) and 72 hours (at E16.5) (B) post-injection, after unilateral injection of AAV-PHP.eB at E13.5.

**Fig S5.**
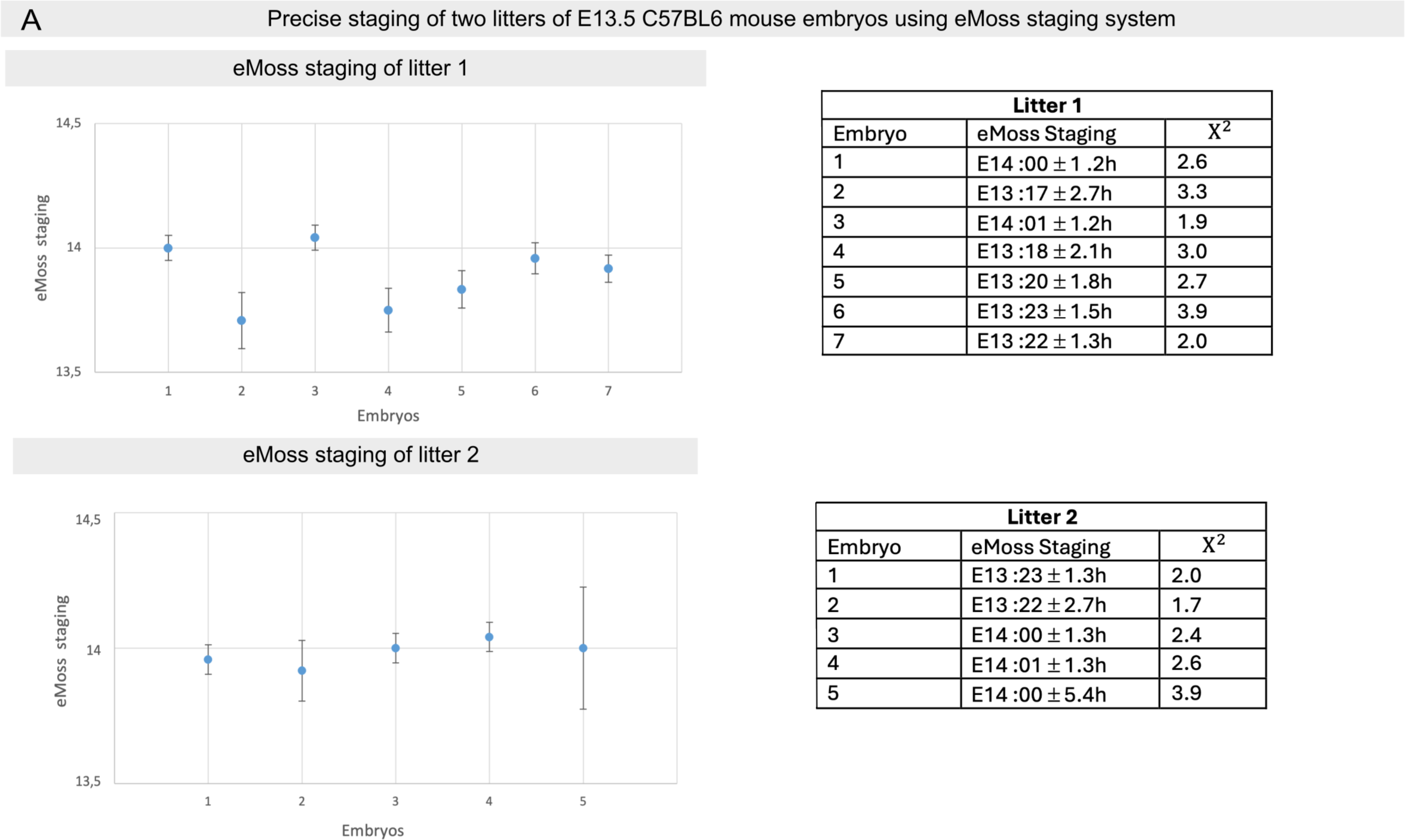
Precise staging of two litters of E13.5 C57BL6 mouse embryos using eMoss. Graphical representation of eMoss staging results using hindlimbs of two litters of mouse embryos staged E13.5 based on the day when the vaginal plug was observed. Only the results for which the values were bellow 5 were considered as reliable and were used for the analysis [20]. Absolute values generated by eMoss and used to built the graphs for litters 1 and 2 are in the respective tables.

## Notes

https://www.protocols.io/view/in-utero-inner-ear-aav-injection-kqdg3x62pg25/v1

### Competing Interest Statement

The authors have declared no competing interest.

