## Supplementary material for "*In utero* adeno-associated virus (AAV)-mediated gene delivery targeting sensory and supporting cells in the embryonic mouse inner ear": https://www.protocols.io/view/in-utero-inner-ear-aav-injection-kqdg3x62pg25/v1

Raphaë

Raphaël Etournay

**Protocol Info:** Carla Maria Barbosa Spinola, jacques.boutet-de-monvel, saaid.safieddine, ghizlene.lahlou, Raphaël Etournay . In utero inner ear AAV injection . **protocols.io**
https://protocols.io/view/in-utero-inner-ear-aav-injection-cw4kxguw

**Created:** Jul 11, 2023

**Last Modified:** Mar 13, 2024

**PROTOCOL integer ID:** 84844

**Keywords:** Carla Barbosa Spinola

**Funders Acknowledgement:**
Fondation pour l'Audition
Grant ID: FPA-IDA-STARTING-GRANT
Agence Nationale de la Recherche
Grant ID: ANR-21-CE13-0038-013-01
Agence Nationale de la Recherche
Grant ID: LabEx LIFESENSES ANR-10-LABX-65

ABSTRACT

In vivo gene delivery to tissues using adeno-associated vector (AAVs) has revolutionized the field of gene therapy. Yet, while sensorineural hearing loss is one of the most common sensory disorders worldwide, gene therapy applied to the human inner ear is still in its infancy. Recent advances in the development recombinant AAVs have significantly improved their cell tropism and transduction efficiency across diverse inner ear cell types to a level that renders this tool valuable for conditionally manipulating gene expression in the context of developmental biology studies of the mouse inner ear. Here, we describe a protocol for *in utero* micro-injection of AAVs into the embryonic inner ear, using the AAV-PHP.eB and AAV-DJ serotypes that respectively target the sensory hair cells and the supporting cells of the auditory sensory epithelium. We also aimed to standardize procedures for imaging acquisition and image analysis to foster research reproducibility and allowing accurate comparisons between studies. We find that AAV-PHP.eB and AAV-DJ provide efficient and reliable tools for conditional gene expression in cochlear sensory and supporting cells, from late embryonic stages on, in the mouse inner ear.
The protocol is caried out in accordance with the European Directive 2010/63/EU of September 22, 2010 on the protection of animals used for scientific purposes. These methods cause neither distress nor pain to the animals.

MATERIALS

**Surgical Material**

| Material | Name | Company | Reference |
|---|---|---|---|
| Automatic Injector | Injector Nanoliter 2020 and Micro T2 controller | WPI | NANOLITER2020 |
| | Nanoliter Injector Footswitch | | 13142 |
| Capillary | Glass capillaries (300) - 1.14 mm OD | WPI | 504949 |
| | Mineral Oil | FertiPro | MT276 |
| | Backfilling syringes | WPI | MF34G-5 |
| Capillary Puller | P-97 Micropipette Puller | Sutter Instruments | |
| Laparotomy and suturing | Needle driver | WPI | 14109 |
| | Graefe Forceps | FST | 11050-10 |
| | Suture Novosyn (5/0) 45cm DR12 | B.Braun Medical | C0068134 |
| | Ball-tipped scissors | FST | 14109-09 |
| | Gauzes | | |
| | Sodium Chloride 0.9% | Lavoisier | 3400930578452 |
| | Fine Scissors | FST | 15000-08 |
| Pre-operatory preparation | Ocry-Gel (10g) | TVM | 03700454505621 |
| | Hair Clipper | Wella | Contura Chrome 4015600113261, hair clipper HS61 |
| | Vetedine Solution | Vetoquinol | 03605870001385 |
| | Gauzes | | |
| | Raucodrape (Adhesive drapes) | Lohmann & Rauscher | 25440 |
| | Heat block and heat Pad | | |
| | Bead sterilizer | FST | 18000-45 |
| Analgesia and anesthesia | Vetergesic Multidose 0.3mg/ml (Buprenorphine) | CEVA | 0103411112112685 |
| | Laucaine (Lidocaine) | MSD | 05017363520132 |
| | Isoflurane | Isovet/Isocare | |
| | Compact Anesthesia Module | Minerve | |

Table 1: List of the surgical material used

### Preparation of animals and materials

2d

1 Pregnant female mice must be acclimated to the local animal facility at least 2 days prior to surgery. One day before surgery, weight the mouse, and observe its general behaviour.

2 Decontaminate the materials used for surgery. Either chemical decontamination (hydrogen peroxide) or autoclave, depending on the type of material.

3 Prepare the capillaries using the following micropipette puller program: **heat:**600**, del:**1**, vel:**55

### Preparation for surgery

30m

4 Pre-warm at least **50ml of a sterile 9%NaCl solution**, in a dry bath heatblock set at 37 °C and tur on the heat sterilizer

5 Weight the dam, at least 00:30:00 before the surgery, and administer the appropriate dose of analgesia (**Vetergesic 0.1mg/kg**) by a subcutaneous injection of the drug volume into the lose over the dam's back scruff. 30m

***Volume to be administered (ml) = [dose** (mg/kg) x **weight** (g)] / [**Buprenorphine** (mg/ml) x **1000]***

Note

*Make sure not to inject more than 1ml, dilute the vetergesic in 0.9% NaCl solution.*

During this time prepare the pre-operatory, the operatory and the recovery areas.

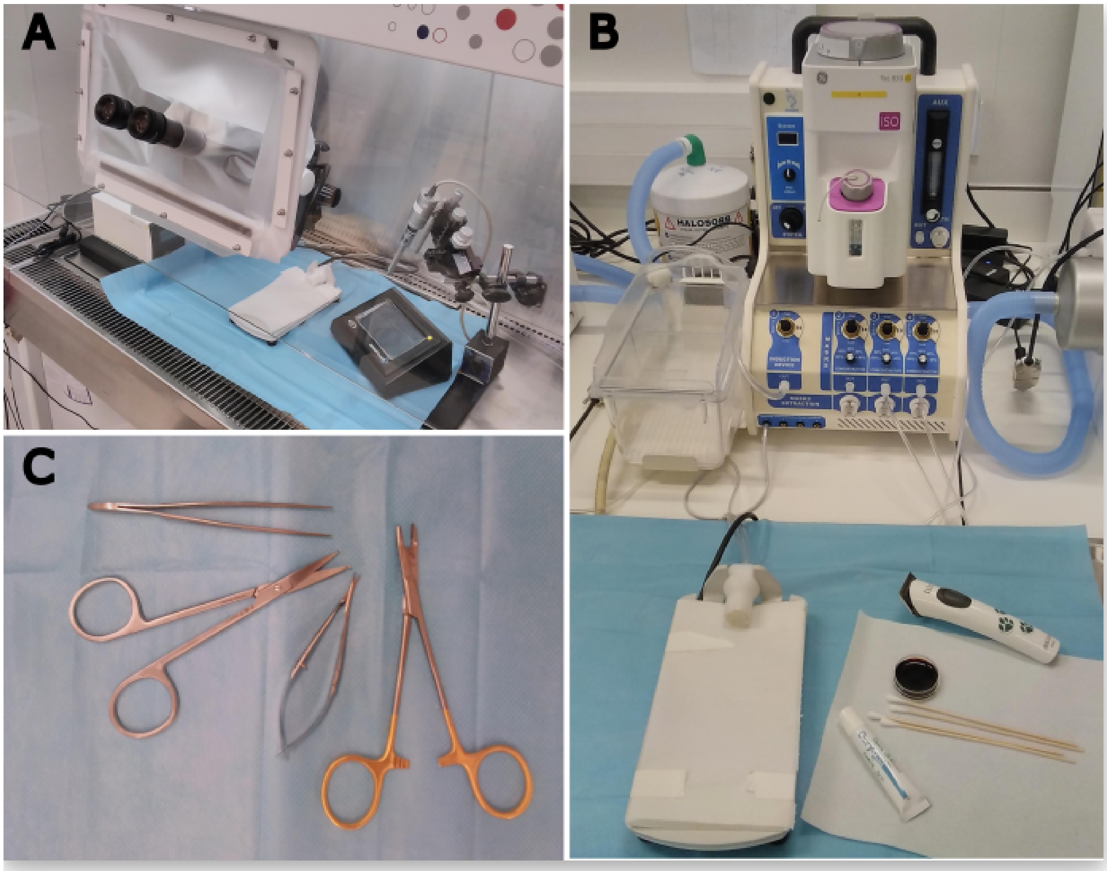


**Figure S3**: Surgery and injection materials
A. Safety cabinet equiped with a binocular in the operatory area. B. The preparatory area equiped with an anesthesia set up. in this are the pregnant dam is shaved in the stomach area after which an antiseptic solution of vetedine is applied. A solution of ocrygel is applied to both eyes. C. Graefe forceps, ball-tipped scisors, fine scisors as well as a needle holder used for the ventral laparotomy, as well as suturing.

6 Turn on the bead sterilizer and prepare the **pre-operatory** by decontaminating the area with Anios SurfaSafe, and Ethanol. Place a gauze over the heat pad where the dam will be placed. Organize the space around the heat pad such that the materials needed for skin antiseptisis and the ocrygel are reachable.

7 Decontaminate the **operatory area**, under the safety cabinet. Clean the materials with Anios SurfaSafe and Ethanol, place a gauze over the heat pad where the dam will be placed for surgery and install the nano-injector equipement.

7.1 Under the safety cabinet, **pre-fill one of the pulled-glass capilaries** with mineral oil, using the backfilling syringe. Fill the capillary up to 75% of its capacity and avoid creating bubbles by placing the tip of the fine srynge the closest to the capilary tip. Cut the tip of the pulled capillary, and make sure that the tip is around 0.7 cm long. Install the capillary in the nanoinjector, and use the micro-T2 controller to inject as much mineral oil as possible, from the prefilled capillary.

### Surgical procedure

1h

8 Anesthetize the dam in a induction chamber with 4% isoflurane, at 2L/min rate. Remove the animal from the chamber and place it in prone position over the heat pad, placing its snout inside the a cone shaped isoflurane mask. Adjust the isoflurane to 2% at 0.6L/min rate.

9 Apply the eye lubricant (ocrygel) to both eyes. Turn the mouse to supine position and shave the fur from the abdominal part with the hair clipper. Then scrub the skin with vetedine at least three times. Quickly transfer the dam to the surgery area, and install the animal to the heat pad connected to an isoflurane mask under the safety cabinet in the surgery area.

10 Place a racodrape over the female such that only the skin area, to be incised is exposed. Check that the mouse is fully anesthetized by pinching the tail.

11 Perform a first incision of the abdominal skin of about 1.5 cm, to expose the abdominal wall using a midline aproach. Irrigate the peritoneum with 0.3 ml/25g of lidocaine 2% (5 mg/kg). Expose the abdominal cavity by an incision through the linea alba of about 1.5 cm. Place at least two gauzes soaked in warm 0.9% NaCl to each side of the incision.

**Note**

For the laparotomy, the incision can be made by pulling the stomach skin with the graefe forceps and making a frist small incision in the skin using the fine scissors, and then you can enlarge the incision by using the balll tipped scisors. In othher hand the incision can also be perfoormed using a scalpel.

12 Carefully squeeze out the the uterine horns by pushing the sides of the belly by hand and lay them the over the soaked gauzes. Recurrently hydrate the uterine horns with sterile warm 0.9% NaCl solution thoughout the procedure.

13 Fill the capillary with at around 3.5 µl of viral solution, enough for at least 2 consecutive injections.

14 Take the embryos one by one, and turn them to one side such that one of the two ears is facing up. The relief of the external ear is clearly visible under the binocular magnifying glass, from E13.5 and later and visualization its aided by trans-illumination of the uterus, with a fiber optic light. The ear is located at the center of a square traced by the cephalic vein in front and its anterior and posterior branches above and bellow. ear

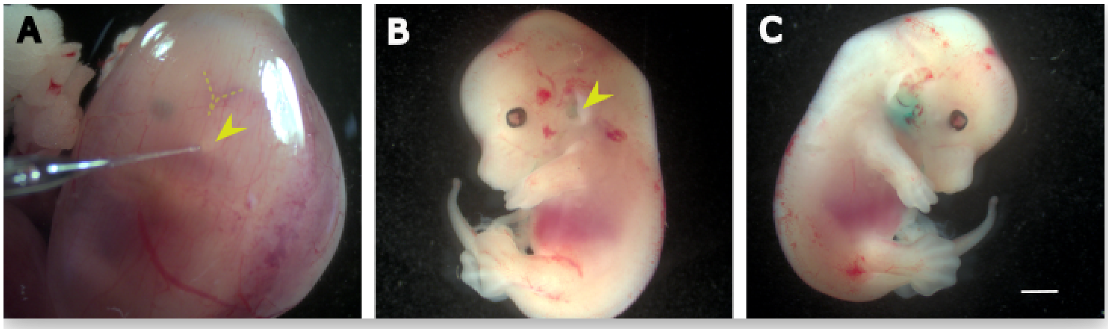


**S2: Inner ear diffusion of FastGreen administered at embryonic stage**
In A, a photo of an embryo inside the uterine horn.
Notice the capillary pointing towards the external ear, located bellow the 'V'-shaped veins corresponding to the transverse and sigmoid sinus veins B-C) Micrographs of E13 mouse embryos after the inner ears were injected with FastGreen; B) the injected (left) ear; C) the contralateral ear, which shows a higher concentration of dye. Scale bar, 1mm.

15 Insert the capillary inside the inner ear region, as perpendicular to the head's sagittal plane as possible.

Note

Avoid inserting the capilary near blod vessels, as there is a risk of hemorrage and embryo lethality. Slightly move the capillary once inside the embryo's head to make sure it is well positioned inside.

16 Trigger the automatic delivery of 1.2 µl of viral solution at constant rate of 600 nl/min. Repeat the procedure for the next embryos, and refill the capillary with with the viral solution, between each two or three consecutive injections.

17 Once all embryos are injected, place the uterine horns back into the dam's abdominal cavity. Irrigate the cavity with warm 0.9% NaCl solution.

18 Close the peritoneum layer by doing a continuous suture, using a resorbable 5-0 Vicryl suture. Then suture the skin layer by single interrupted stitches, using non-resorbable suture Dacron 6-0.

### Post-operatory recovery

19 Place the dam in supine position inside a cage, over a heating pad, until awakening. Cover the floor of the cage with cottons, and jelly recovery food (DietGel), as well as normal diet food.

Note

If the dam shows any signs of pain provide a seconde analgesic dose or Vétérgesic.

20 Monitor the animals for the next few days until full recovery, at least once a day. Inspect for any sign of illness or pain, and provide analgesic solution (Vetergesic (0.1mg/kg)) for the next two days after surgery.

### Collecting inner ears of AAV injected pups

21 Euthanize the newborn pups at P0, by decapitation, and collect the inner ears. Euthanize the dam by exposing it to increasing CO2 concentration using a dedicated C02 box, for at least 10 minutes.
